# A cryo-ET survey of intracellular compartments within mammalian axons

**DOI:** 10.1101/2021.03.29.437454

**Authors:** H E Foster, C Ventura Santos, A P Carter

**Affiliations:** Structural Studies Division, MRC Laboratory of Molecular Biology, Cambridge, United Kingdom; Max Planck Institute of Molecular Cell Biology and Genetics, Dresden, Germany

**Keywords:** Cytoskeleton, Cryo-ET, Neuroscience, Trafficking, Organelles

## Abstract

The neuronal axon contains many intracellular compartments which travel between the cell body and axon tip. The nature of these cargos and the complex axonal environment through which they traverse is unclear. Here, we describe the internal components of mammalian sensory axons using cryo-electron tomography. We show that axonal endoplasmic reticulum has thin, beaded appearance and is tethered to microtubules at multiple sites. The tethers are elongated, ∼ 7 nm long proteins which cluster in small groups. We survey the different membrane-bound cargos in axons, quantify their abundance and describe novel internal features including granules and broken membranes. We observe connecting density between membranes and microtubules which may correspond to motor proteins. In addition to membrane-bound organelles, we detect numerous proteinaceous compartments, including vaults and previously undescribed virus-like capsid particles. The abundance of these compartments suggests they undergo trafficking in axons. Our observations outline the physical characteristics of axonal cargo and provide a platform for identification of their constituents.

## Introduction

Axons are slender processes which extend from the neuronal cell body and transmit electrical signals to neighboring cells. Proper axonal function requires a constant supply of components to the synaptic termini. Since mature axons contain few ribosomes (Bartlett and Banker, 1984; Court et al., 2008), the majority of their building blocks are transported from the cell body. This intracellular transport relies on the microtubule cytoskeleton, which lines and shapes the axon.

Delivery of components to the axon tip (i.e. anterograde transport) relies on the kinesin family of motors. The speed the of transport can be slow or fast depending on the cargo. Cytosolic proteins including many cytoskeletal elements are transported slowly, at rates of 0.2-10 mm/day (reviewed in Roy (2014)). In contrast, membrane organelles and associated components can travel up to 20 times faster, at rates of 50-200 mm/day. This fast transport quickly delivers Golgi-derived vesicles containing neuropeptides (Barkus et al., 2008), transmembrane channels (Akin et al., 2019) and receptors (Gumy et al., 2017; Zhao et al., 2011) into the axon. The return of components back to the cell body (i.e. retrograde transport) is exclusively fast (Vallee and Bloom, 1991). These cargos include endocytic vesicles, autophagosomes and lysosomes which rely on the cytoplasmic dynein-1 motor (Gowrishankar et al., 2017; Maday et al., 2012; Olenick et al., 2016). This retrograde transport is important for the transmission of growth factor signals and the clearance of damaged components (Cai et al., 2010; Chakrabarty et al., 2018). Axons also contain mitochondria (Saxton and Hollenbeck, 2012) and a network of smooth endoplasmic reticulum (ER) (Wu et al., 2017). These are spread throughout axonal processes to provide ATP and lipids, in addition to regulating calcium homeostasis. Overall, the axon shaft is highly dynamic, with different components navigating their way through the axoplasm at different rates.

Cryo-electron tomography (cryo-ET) is becoming an invaluable tool to understand cellular ultrastructure (Albert et al., 2019; Allegretti et al., 2020; Bykov et al., 2017; Jordan et al., 2018), and can provide sub-nanometer resolution structural information *in situ* (Himes and Zhang, 2018; O’Reilly et al., 2020; von Kügelgen et al., 2020). To better understand the architecture of intracellular compartments in the axon shaft, we imaged mammalian dorsal root ganglion (DRG) neurons using cryo-ET. We show that the ER has a thin, beaded appearance and discover that it is connected to the microtubule cytoskeleton by collections of short tethers. Unexpectedly, we find that some small vesicles and late endosomes contain condensates in their lumen. On the surface of small vesicles, we identify large protein complexes and in some cases identify connections to microtubules. In addition to the membrane-bound components, we detect many virus-like capsid structures, suggesting proteinaceous shells may be frequently transported within axons. Our work provides insight into the nature of axonal compartments and how they interact with their surroundings.

## Results and Discussion

### Cytoskeletal components observed in axons

DRG neurons transmit information from the periphery to the spinal cord. *In vitro* cultures have been widely used to study microtubule-based cargo transport (Akin et al., 2019; Cui et al., 2007; Delcroix et al., 2003; Gumy et al., 2017; Maday and Holzbaur, 2014). A key advantage of this system is that all of the processes which extend from the cell body are axon-like (Riederer and Barakat-Walter, 1992), with uniform microtubule polarity (Kleele et al., 2014). We plated adult mouse DRG neurons on electron microscopy (EM) grids and found axonal processes extended over the grid surface within 4 days. We labelled acidic vesicles using LysoTracker and performed live imaging. These vesicles moved bidirectionally (Figure 1A,B), indicating that neurons grown on EM grids are capable of axonal transport.

**Figure 1.**
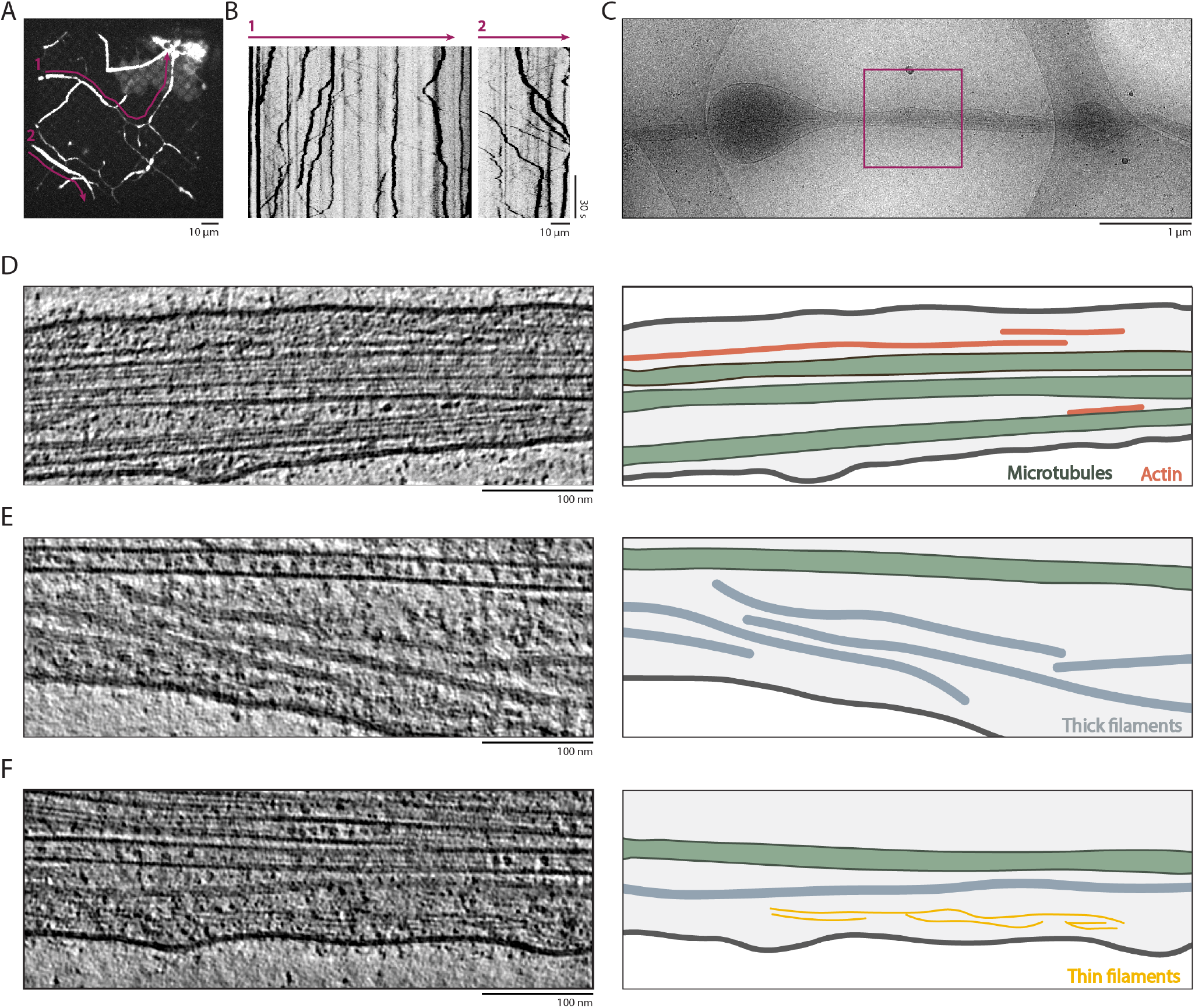
Filaments within DRG neurons grown on EM grids. A) Maximum intensity projection of a time-lapse movie of DRG neurons labelled with LysoTracker Far Red dye, showing the axon distribution across the grid surface. Purple lines indicate the position of axons used for kymograph generation. B) Kymographs generated from axons indicated in A showing bidirectional transport of LysoTracker-positive (acidic) vesicles. In these images, the position of the cell bodies with respect to the axons, and therefore the direction of transport, cannot be determined. C) Overview transmission electron microscopy image of a thin axon targeted for cryo-ET acquisition. Purple box indicates a region targeted for data acquisition. D-F) Slices through tomograms showing filaments within axons. Right hand panels show tracing of plasma membrane (dark grey), microtubules (green), actin (orange), ’thick’ filaments (blue) and ’thin’ filaments (yellow).

We acquired two cryo-ET datasets of DRG neurons at sites where they grow across the holes in the carbon foil of the EM grids (Figure 1C). One dataset of 30 tomograms included both thick and thin regions of axons, allowing us to survey both large and small organelles. Tomograms that were thicker than 300 nm had lower contrast but the characteristic features of the vesicles were clearly visible. Our second dataset focused on regions thinner than 200 nm. All 39 tomograms in this dataset were of sufficient quality to follow the paths of filaments including microtubules. Larger membrane-bound organelles were absent in from thin regions and are therefore not found in the second dataset. In this paper, we survey the ultrastructure of organelles using the first dataset and the second to provide fine structural details.

We observed four distinct classes of filaments in the axoplasm. The first were microtubules (Figure 1D-F), which we observed in all but two of our tomograms. We provide a more detailed analysis of these microtubules in a companion manuscript (Foster et al., 2021). The second class of filaments was actin (Figure 1D) which we identified based on their (1) characteristic “zig-zag” appearance, (2) diameter of 6.1±1.1 nm, and (3) 6 nm layer lines in Fourier transforms (Figure 1D, S1A and S1B) (Chou and Pollard, 2019; Hanson and Lowy, 1963; Narita et al., 2012). We observed actin filaments distributed in most (48/62) of the tomograms. While microtubules typically extended beyond the imaged volume, actin filaments were short, with an average length of 250 nm (Figure S1C). Similar to a recent cryo-ET study of human organoid axons (Hoffmann et al., 2020), the observed actin filaments ran roughly parallel to the microtubule network rather than forming rings around the axon shaft. This actin may correspond to “deep” actin which is dynamic and is transported slowly through axons (Chakrabarty et al., 2018; Ganguly et al., 2015).

The identify of the other filament classes were less clear. One had a diameter of 8.5 nm (Figure 1E and S1A) and typically ran the full length of the imaged volume, suggesting they are longer than our 1.2 µm field of view. We observed 83 of these ’thick’ filaments in 41 out of the 62 tomograms. Their diameter is similar to purified neurofilaments (Wagner et al., 2004) and to non-neuronal intermediate filaments previously observed by cryo-ET (Chakraborty et al., 2020; Mahamid et al., 2016). The major intermediate filament components in DRG neurons include the three neurofilament isoforms (NF-H, NF-M and NF-L) and peripherin (Fornaro et al., 2008). These ’thick’ filaments are therefore likely intermediate filaments.

Current models suggest intermediate filaments are built from coiled-coil dimers which oligomerize into bundles called unit length filaments (ULFs). The ULFs are 60 nm long and anneal end-on-end to form long polymers (Herrmann and Aebi, 2016). Unlike actin, the ’thick’ filaments in our tomograms showed no clear reflections in a Fourier transform (Figure S1D). This suggests they are heterogenous in composition or the repeating unit is difficult to detect in Fourier transforms. We performed subtomogram averaging and found they are tubular, with less density in their core (Figure S1E). This hollow arrangement is different from models of intermediate filament bundle formation (Herrmann and Aebi, 2016).

The final class of filaments are 3 nm in diameter and shorter than the others, with an average length of 116 nm (Figure S1A,C). These ’thin’ filaments typically ran parallel to the microtubules and sometimes appeared branched or in groups (Figure 1F and S1F). Their diameter suggests that they may be coiled-coils. A possible candidate is spectrin, which binds the plasma membrane and actin to form the membrane-associated periodic skeleton (He et al., 2016; Xu et al., 2012), and is transported within axons (Levine and Willard, 1981; Lorenzo et al., 2019). Alternatively, they could be intermediate filament subunits which are yet to assemble.

In summary, we observed that microtubules, actin, intermediate and ’thin’ filaments form a loosely packed parallel array. This is distinct from the meshwork previously reported to form the membrane-associated periodic skeleton (He et al., 2016). Together with the ER network we describe below, these components form the environment that must be navigated by membrane-bound organelles during axonal transport.

### The ER is connected to the microtubules through short tethers

Smooth ER extends from the cell body to form a network of thin tubules in axons (Terasaki, 2018; Wu et al., 2017). It is a key site for lipid biosynthesis and distribution (de Chaves et al., 1995; Gould et al., 1983; Vance et al., 1991), an effector of calcium homeostasis (de Juan-Sanz et al., 2017; McGraw et al., 1980) and a source of autophagy membranes (Maday and Holzbaur, 2014). In 35 of 69 tomograms, the ER was clearly visible and continuous, with regions of narrow tubules as well as more open stretches (Figure 2A,B). The narrow tubules were present in 30 tomograms and had undulating shape, where the two lipid bilayers come in contact. This “beaded” morphology is also reported in a recent study of human axons (Hoffmann et al., 2020). In 8 tomograms we found electron dense deposits in the ER lumen (Figure S2A). These appear similar to calcium deposits within mitochondria (Wolf et al., 2017). Their presence has not been described in recent cryo-ET studies, but the ER in neurons is known to sequester calcium and calcium-containing sites were previously observed in the ER of presynaptic nerve terminals (McGraw et al., 1980).

**Figure 2.**
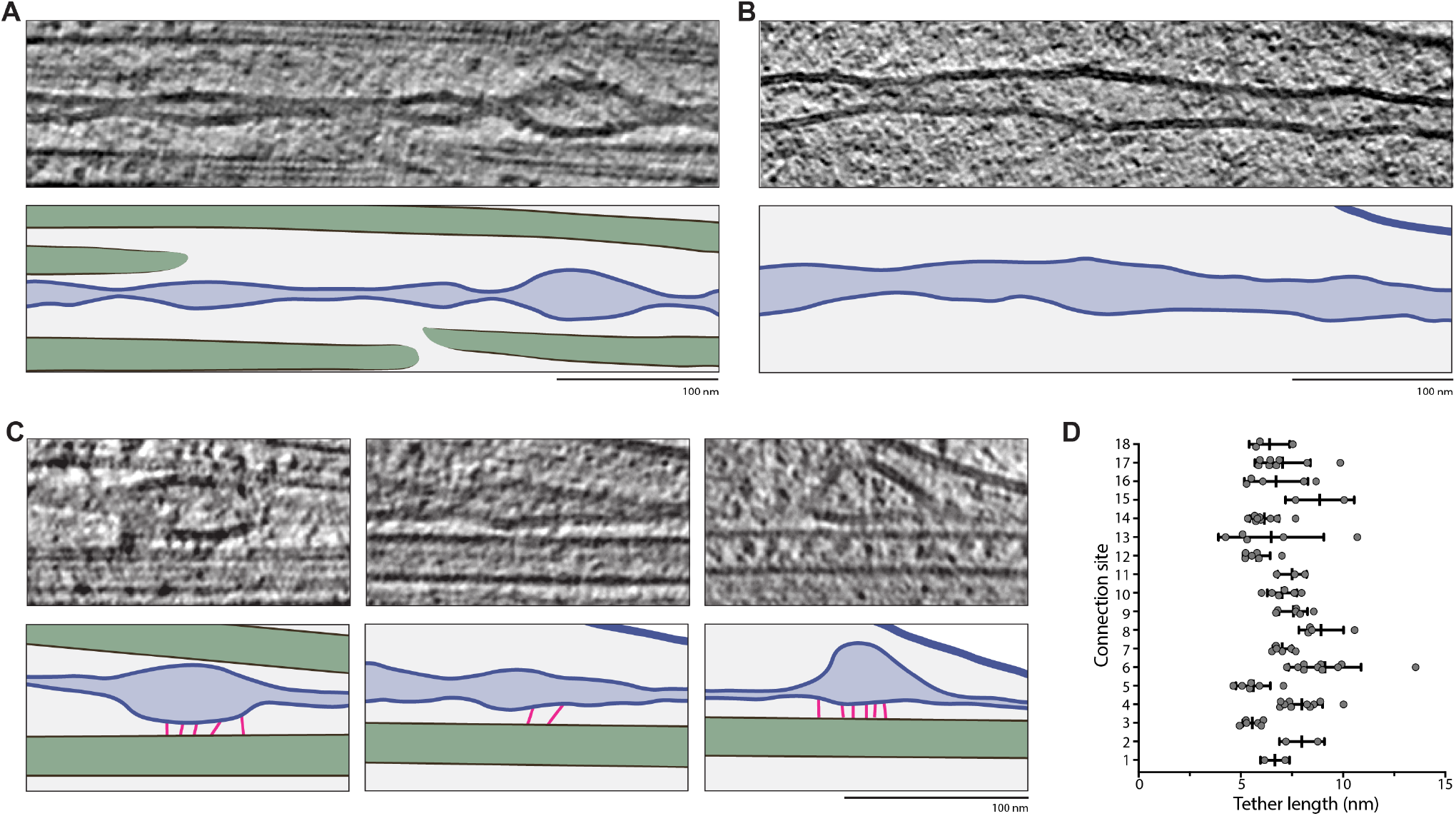
ER-microtubule contact sites within axons. A-C) Slices through tomograms showing ER morphology. In the cartoon below each tomogram, microtubules are green, ER membrane is blue, ER lumen is light blue and connections between ER and microtubules are pink. A) shows narrow ER with beaded morphology. B) depicts an ER section with wider lumen and more open morphology. C) shows regions with a series of short, regular connections linking the ER and microtubules. D) Quantification of the length of tethers between ER and microtubules at each connection site.

A key question about ER morphology in axons is how it is stretched and maintained. In our data, we identified clear connections between microtubules and the ER (Figure 2C). We found 18 contact sites within 12 of 69 tomograms (Figure S2B). They predominantly lie in regions of beaded ER with a wider lumen. An average site contains 5 short tethers, each being 7±2 nm long (Figure 2D). As they are small, it is likely there are more connection sites that we were unable to detect. The prevalence of the tethers suggests they play a role in stabilizing the ER and maintaining its structure.

The average length of the tethers is consistent with a ∼ 50 amino acid coiled coil. This is different from the dimensions of the microtubule motors responsible for ER distribution in axons (Farías et al., 2019). They are too short to correspond the kinesin motors (Hirokawa, 1998) and have smaller dimensions than dynein/dynactin complexes (Grotjahn et al., 2018; Urnavicius et al., 2018, 2015). In addition to motors, a number of ER-resident transmembrane proteins have been reported to directly bind the microtubule lattice. These include p180, CLIMP-63, REEP-1 and Sec61β (Klopfenstein et al., 1998; Ogawa-Goto et al., 2007; Park et al., 2010; Zhu et al., 2018). CLIMP-63 localizes to rough ER and is therefore mainly excluded from axons (Cui-Wang et al., 2012; Farías et al., 2019). p180 has been shown to stabilize the ER and microtubule networks in axons (Farías et al., 2019) but is likely too big to correspond to the tethers we observe (Figure S1C). REEP-1 and Sec61β are possible candidates but lack any coiled coil prediction (Figure S2C). It is therefore likely the tethers are formed from an as-yet-unidentified factor.

### The morphology of single membrane cargos

In the dataset containing both thin and thick axonal regions, we identified 247 membrane-bound compartments across 30 µm of axon length. 202 of these were single membrane vesicles, whereas 45 compartments contained more than one lipid bilayer. Of the single membrane vesicles, we saw examples with both light- and dark-lumen (Figure 3A,B and S3A-C). The majority of light-lumen vesicles had a diameter of 40 - 60 nm which is similar to the size of synaptic vesicles frequently observed in the presynapse of hippocampal neurons (Tao et al., 2018). We also detected larger light-lumen vesicles. These had a diameter of 70 - 170 nm and may correspond to less well characterized axonal compartments such as synaptic vesicle precursors or early endosomes (Pace et al., 2020; Wang et al., 2016; Wu et al., 2017). Vesicles with a dark lumen had an average diameter of 95 nm with the majority ranging from 70 to 120 nm. These are likely dense core vesicles (DCVs) which have diameters between 70 and 100 nm (Persoon et al., 2018; Sorra et al., 2006; Tao et al., 2018; Zhai et al., 2001). Previous studies have shown that DCVs are distributed throughout hippocampal axons and are transported to presynaptic membranes where they release the neuropeptides or neurotrophins stored in their lumen (Bost et al., 2017; Xia et al., 2008). They can also carry scaffold proteins which are deposited at the presynapse to form the ‘active zone’ (Shapira et al., 2003; Zhai et al., 2001).

**Figure 3.**
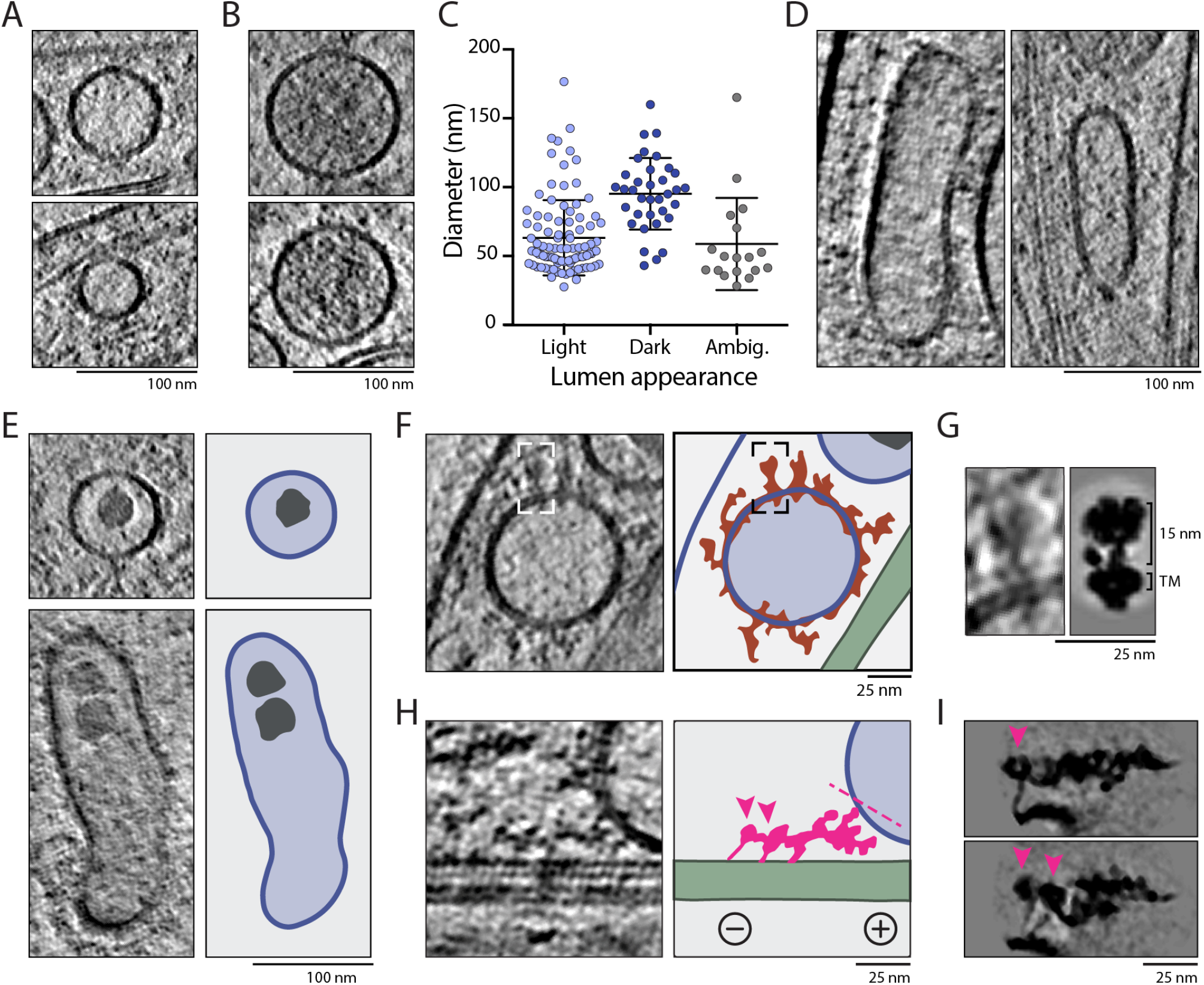
The fine structure of uni-lamellar vesicles within axons. (A,B) Tomogram slices showing two example light-lumen (A) or dark-lumen vesicles (B). C) Distribution of vesicle diameter showing spherical light-lumen vesicles are smaller on average (63.4±27.4 nm, n=98) than spherical dark-lumen vesicles (95.4±26.0 nm, n=35). Those with unclear lumen density are classed as ambiguous (59.0±33.5 nm, n=18). D) Slices through tomograms showing typical morphology of elongated vesicles. E) Examples of spherical and elongated vesicles containing non-membrane bound granules. Cartoon shows membranes (blue), vesicle lumen (light blue) and smooth sided granules (grey). F) Slice through a tomogram showing large membrane-bound proteins visible on the vesicle surface. Cartoon is colored as in E, with protein density outlined in orange and microtubules in green. G) An enlarged view of the protein densities boxed in F, which show similarity to V-ATPase. The right panel shows a low-pass filtered projection of the V-ATPase (EMD-31217) to scale. Position of transmembrane helices (TM) are indicated. H) Slice through a tomogram showing motor-like connections between the vesicle and microtubule. In the cartoon, connecting density is pink. Arrow heads indicate globular density which may correspond to dynein motor domains and dashed line indicates approximate length of dynactin filament (38 nm). Plus and minus indicate microtubule polarity. I) Low-pass filtered projection of the dynein/dynactin complex (EMD-7000) using two different projection angles to show the possible appearance of motor complexes on microtubules.

We noticed a subset (47/202) of the light- and dark-lumen vesicles were non-spherical and appeared elongated (Figure 3D, S3D). In our data, the elongated compartments were observed in the same tomograms as spherical vesicles and found in both wide and narrow axons, suggesting external force is not responsible for their shape. Instead, it is possible that the lipid and protein composition of the membrane is responsible for their shape(McMahon and Gallop, 2005). Similar tubulovesicular structures were previously reported to undergo anterograde transport in axons (Hirokawa et al., 1991; Tsukita and Ishikawa, 1980).

A subset (6/202) of single-membrane vesicles contained granules in their lumen (Figure 3E). These granules had smooth sides and were not surrounded by a lipid bilayer, suggesting they are biomolecular condensates. We noticed that the granules were non-spherical with angular corners. We saw some granules in close proximity to each other (Figure 3E), suggesting they do not readily coalesce and so have gel-like rather than liquid-like properties (Alberti et al., 2019). While there has been intense study of phase separated particles in the cytoplasm, they have not been previously seen in membrane-bound compartments. Although it is not clear what these granules are made of, there is indirect evidence that small molecules such as ATP and choline can be stored in a gel-like matrix in neuronal vesicles (Rahamimoff and Fernandez, 1997; Reigada et al., 2003). Alternatively, these vesicles may be DCVs which have undergone piecemeal degranulation, a process where the internal cargo is reduced and only occupies a small space inside the vesicle lumen. This process has been described for mast cell granules (Dvorak, 2005; Melo and Weller, 2010) and is also proposed to occur in hippocampal neurons (Crivellato et al., 2005).

### Protein decorations on membrane-bound organelles

Using current cryo-ET methods, *in situ* identification of membranes, large macromolecules such as ribosomes, and proteins organized in arrays is achievable (Bykov et al., 2017; Mahamid et al., 2016; Wietrzynski et al., 2020). However, identifying smaller, individual protein complexes is more challenging due to the crowded nature of the cytoplasm and the lack of contrast in cryo-ET images caused by specimen thickness. In our thinnest, best aligned tomograms we observed individual densities protruding from the vesicle surface (Figure 3F). Although many are not easily identifiable, a subset resembled projections of V-ATPases (Abbas et al., 2020) (Figure 3G). These ATP-driven protein pumps are known to be present on the surface of synaptic vesicles, endosomes and lysosomes where they control the luminal pH (Lafourcade et al., 2008; Ohkuma et al., 1982; Overly et al., 1995; Takamori et al., 2006).

Membrane vesicles can be transported through the axon by microtubule motors. Cargo travelling towards the microtubule minus end are carried by cytoplasmic dynein-1, which binds its cofactor dynactin to form a 2.4-3.8 MDa complex which is 65 nm long and 30 nm wide (Grotjahn et al., 2018; Urnavicius et al., 2018, 2015). There are many outstanding questions regarding the mechanism of cargo transport, including how many dyneins are required to power the transport of vesicular cargo. As dynein/dynactin complexes are similar in size to ribosomes and contain a 38 nm actin-like filament, we set out to determine if we can find them within our tomograms. We surveyed regions where membrane vesicles are close to microtubules. We identified 67 objects connecting the membrane to microtubules (Figure S3E). In one of our best tomograms we saw globular ring-like densities attached by thin projections to the microtubule which resemble dynein motors (Figure 3H). These were connected by a ∼ 40 nm tether to the vesicle surface which may correspond to a dynactin filament. This example has similar dimensions and appearance to projections of cryo-EM structures of dynein/dynactin (Figure 3I) (Grotjahn et al., 2018; Urnavicius et al., 2018, 2015).

We determined the direction of individual microtubules within our data, as described in the companion paper (Foster et al. 2021), and used this to find the orientation of each of our putative dynein densities. We found 52% (32/67) of these connections were oriented to the minus end and 48% (35/67) towards the plus end. Actively moving dyneins are expected to point towards the minus end (Can et al., 2019; Grotjahn et al., 2018; Imai et al., 2015). Therefore, the plus-end oriented connections we observed may be other motors or non-motor microtubule associated complexes. Dynein and kinesin have been shown to bind the same cargo (Kendrick et al., 2019; meng Fu and Holzbaur, 2013) and have been proposed to engage in a tug-of-war to determine the organelle transport direction (Hancock, 2014). It is therefore also possible the connections we observed are dyneins that are being pulled along on plus-end directed kinesin cargos. Our analysis of individual complexes in the axoplasm shows that identifying densities on the vesicle membrane and connecting cargo to filaments is feasible. However, accurate identification of motor complexes will require precise labelling approaches or subtomogram averaging of many high-quality tomograms containing dynein/dynactin.

### Multimembrane compartments contain granules and linear membrane sheets

In our data, we observed 45 compartments which contained more than one lipid bilayer. These multimembrane compartments were on average larger than the single membrane vesicles (Figure 4A). Based on physical characteristics, including the number, size and shape of internal membranes, we categorized these compartments into four main groups: mitochondria, multivesicular bodies (MVBs), degradative compartments and smaller compartments with unclear identity.

**Figure 4.**
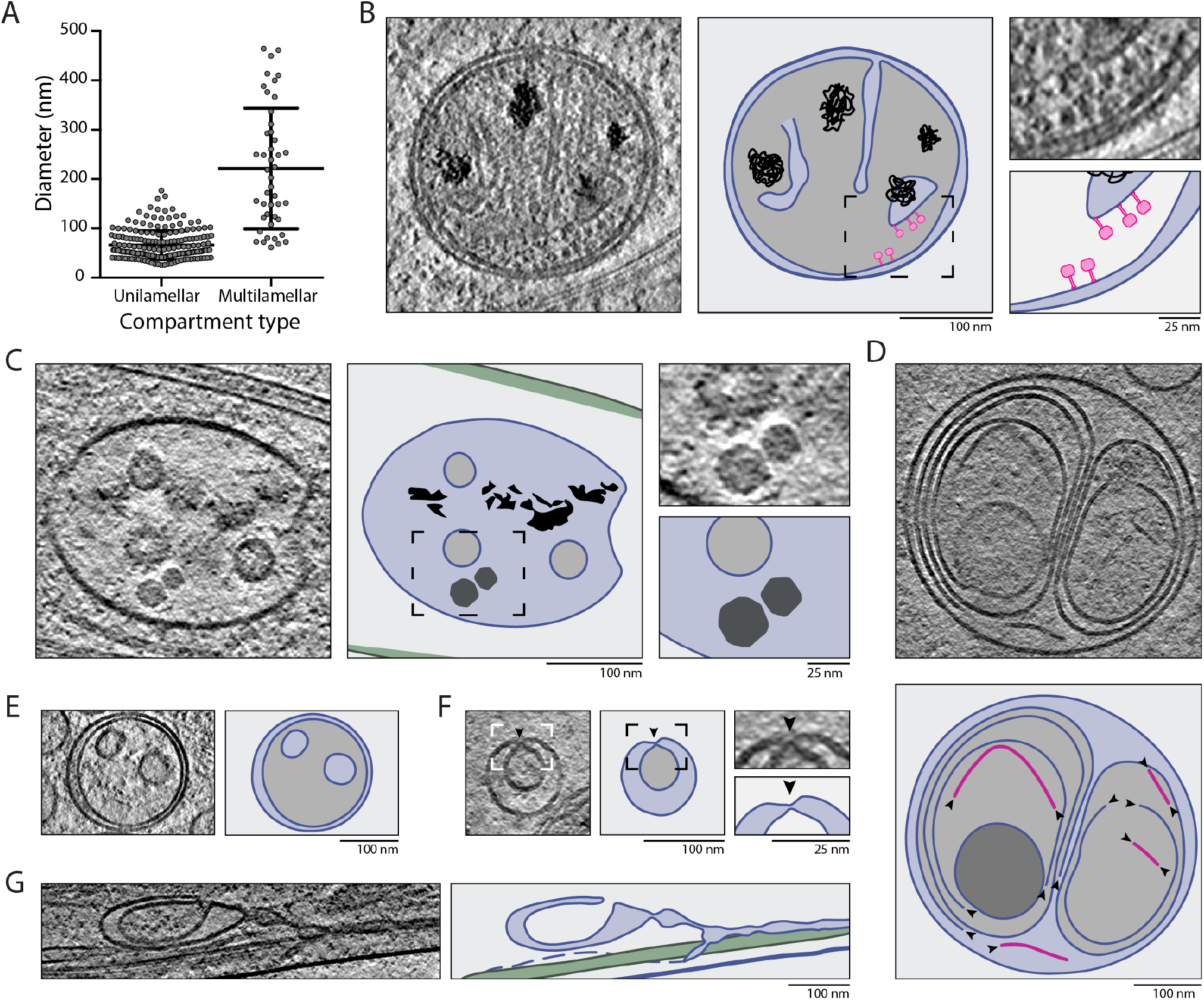
Granules and membrane sheets within multilamellar compartments. A) Measurements of the diameter of membrane-bound compartments within the tomograms showing uni-lamellar vesicles are smaller (66.1±29.1 nm, n=202) than multimembrane vesicles (221.3±122.4 nm, n=45). For elongated compartments, the shortest axis is plotted. B) Slice through tomogram containing a mitochondrion. In the cartoon, membranes are blue, calcium deposits are black and ATP synthase density is pink. Enlarged region shows ATPase heads. C) An example multivesicular body within axons, showing intraluminal vesicles and granules. Cartoon is colored as in B. D,E) Slice through tomogram showing membrane sheets and ruptured vesicles within autophagosomes (D) and endolysosomes (E). In cartoon, membrane sheets are pink. Arrowheads indicate exposed lipid bilayer sides. F,G) Slices through tomograms containing multilamellar compartments of unclear identity. Cartoon is colored as in B. Arrowhead and expanded region in F shows membrane deformation close the outer membrane bilayer.

We identified 9 mitochondria (Kühlbrandt, 2015). These are double membrane compartments with a dark-lumen and a folded inner membrane (Figure 4B, S4A). In some cases, we detected globular density attached to the internal surface of the inner membrane (Figure 4B), which likely correspond to the F_1_F_0_ ATP synthase on the surface of mitochondrial cristae (Ader et al., 2019; Davies et al., 2011). We observed numerous dark deposits in the mitochondrial matrix (Figure 4B, S4A) similar to those observed in the ER (Figure S2A). Mitochondrial deposits have been observed previously and consist of calcium phosphate, which helps regulate intracellular calcium ion concentration (Wolf et al., 2017).

MVBs were identified as compartments with numerous, small vesicles in their lumen (Bartheld and Altick, 2011; Murk et al., 2003). They had a single lipid bilayer surrounding the internal vesicles, all of which were intact (Figure 4C, S4B). MVBs are late endosomes which form through fusion of early endosomes followed by invagination to generate intraluminal vesicles (Klumperman and Raposo, 2014; Murk et al., 2003; Rink et al., 2005). The MVBs we observed in axons are similar to those described in synapses (Schrod et al., 2018) but the novel feature we see are granules in the lumen of 6 out of the 10 compartments (Figure 4C). These granules resemble those present in single-membrane vesicles (Figure 3E).

Another set of multimembrane compartments contained broken vesicles and membrane sheets (Figure 4D). These are likely degradative compartments such as autophagosomes or endolysosomes (Jung et al., 2019; Klumperman and Raposo, 2014; Möbius et al., 2003). Some of the compartments we observed contained many internal membrane layers with an overall dark-lumen (Figure S4C). Others had a more electron transparent lumen and smaller internal vesicles but still contained broken membrane sheets (Figure S4D). Although the link between the identity and physical characteristics of axonal compartments is not well established, we suspect the more dense, complex compartments correspond to autolysosomes. These are formed through encapsulation of cytoplasm including membrane-bound organelles by a double membrane and can therefore contain many membrane layers (Liou et al., 1997; Tammineni et al., 2017; Uemura et al., 2014). We assume the lighter-lumen compartments are endolysosomes, which are formed through fusion of lysosomes with mature MVBs (Luzio et al., 2000).

Previous work has shown the degradative capacity of lysosomes within axons is higher close to the cell body (Cheng et al., 2018; Yap et al., 2018). Our tomograms were collected in thin, distal axons but we still observed broken vesicles and membrane sheets, suggesting these are a common feature of multimembrane compartments. Preliminary membrane breakdown may therefore occur throughout the axon.

The remaining multimembrane compartments lacked broken membranes and contained large internal vesicles, which occupied much of the lumen (Figure 4E-G). 17 out of 45 multimembrane compartments had this morphology, suggesting they are relatively abundant within DRG axons. It is not clear what compartments these correspond to as they have not been widely commented on in previous reports. One contained two small vesicles surrounded by an intact double membrane (Figure 4E), suggesting it is a small autophagosome. Others contained only one vesicle, and in 5 such cases the internal vesicle was puckered close to where it contacted the external lipid bilayer (Figure 4F and S4E), suggesting invagination and membrane scission were taking place. These may be small, immature MVBs or autophagosomes. We also found puckered vesicles within elongated compartments (Figure S4F). In two cases, the vesicles were inside ER tubules (Figure 4G and S4G). The appearance of these internal ER vesicles suggests they are formed through engulfing part of the cytoplasm. The ER in axons is known to be a source of autophagy membranes (Maday and Holzbaur, 2014) and so this sub-group may represent examples of localized small autophagosome generation.

### Axons contain a large number of protein shells that resemble virus-like capsids

So far we have focused on the abundant membrane-bound structures visible in the axoplasm. While cataloguing these observations, we also detected two types of non-membrane bound compartments. The first are vault complexes, highly conserved 13 MDa ribonucleoprotein complexes with well-characterized structure but elusive function (Berger et al., 2008; Kedersha and Rome, 1990; Tanaka et al., 2009; Vasu and Rome, 1995). They have a barrel-shaped central region with caps on either end (Figure 5A, S5A) and are known to be made up of a protein shell (Mikyas et al., 2004). In our tomograms, the density for this shell is clearly thinner than that of adjacent lipid bilayers (Figure S5B). Vaults have been observed in cryo-electron tomograms of various cell types where they co-localize with cytoplasmic structures such as granule beds or autophagy compartments (Carter et al., 2019; Woodward et al., 2015). Previous work has proposed they are transported in axons (Herrmann et al., 1996; Li et al., 1999; Paspalas et al., 2008). We observed that the axonal vaults have similar appearance to those in mammalian epithelial cells or purified from rat liver (Kong et al., 2000; Woodward et al., 2015), with an internal density bound at the inflection between the central barrel and end-caps. We did not observe any co-localization with granules or specific membrane-bound organelles, suggesting they have a distinct function in axons.

**Figure 5.**
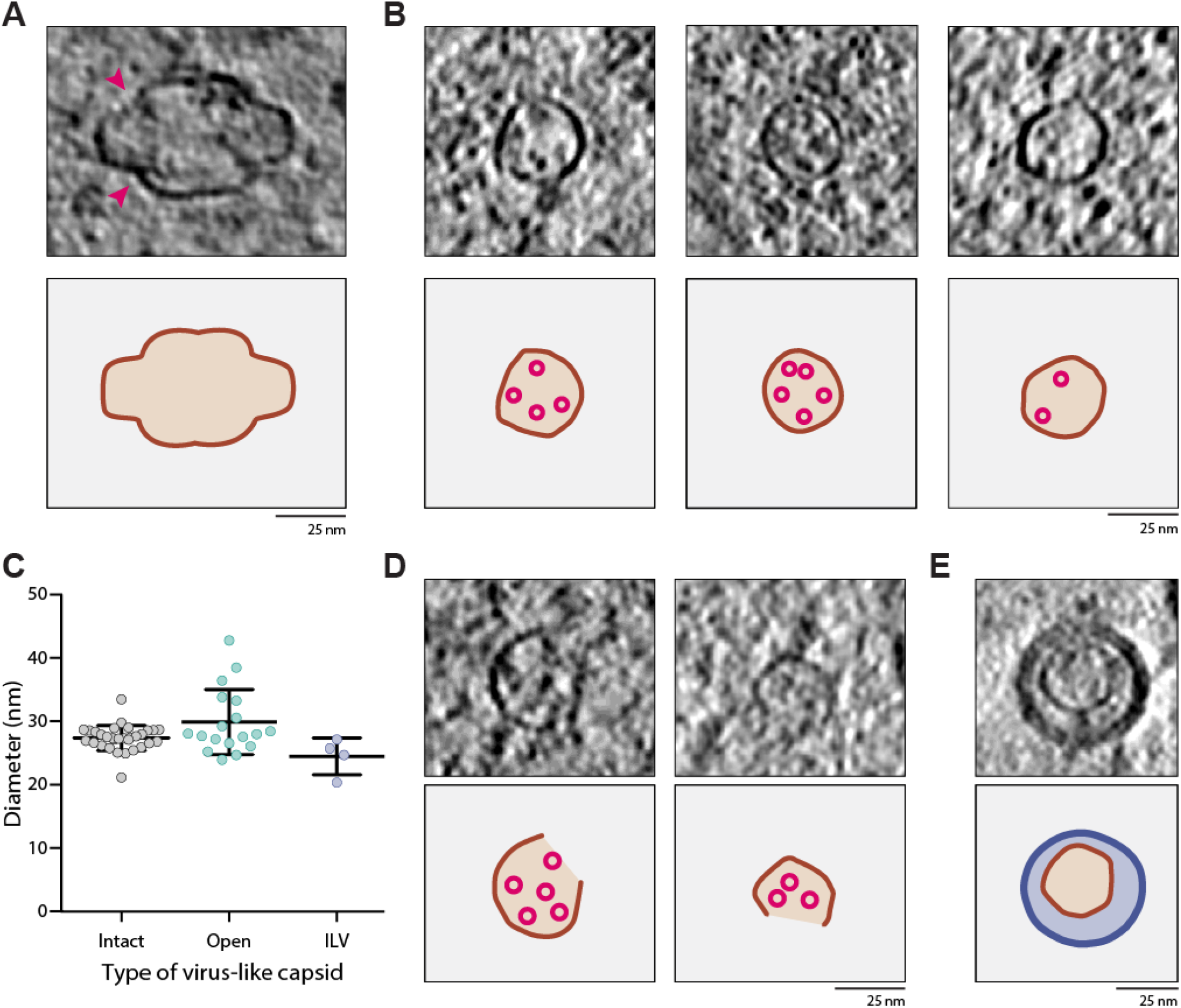
Virus-like capsids within axons. A) Slice through a tomogram showing a typical vault found in the axonal process. Pink arrows indicate internal densities found at the cap/barrel intersection. Lower panel shows outline of vault protein shell in orange. B) Example intact virus-like capsids from within the axonal process. For each example, the lower panel shows the outline of the protein shell in brown and internal densities in pink. C) Scatter plot showing diameter of virus-like capsids classed as intact (27.3±2.0 nm, n=31), open (29.8±5.1 nm, n=18) or within an intraluminal vesicle (ILV, 24.4±3.0 nm, n=4). D) As in B, for open capsid-like particles. E) As in B, for a particle within an intraluminal vesicle of a multivesicular body. In cartoon below, membranes are in dark blue and endosome lumen is light blue.

The second type of non-membrane bound compartment we saw are smaller and roughly spherical (Figure 5B-E and S5C-E). Their wall thickness is similar to that of vaults and they are therefore also likely to be made of a protein shell. We found them in 30 out of the 64 tomograms analyzed, with 53 observed in total. These structures have a diameter of 27±2 nm and angular edges similar to virus capsids (Figure 5B,C). Some of these virus-like capsids are open sided (Figure 5D and S5D), suggesting they can open up in the cytoplasm. In rare cases, they were found within the internal vesicles of MVBs (Figure 5E and S5E) suggesting they can be taken up from the cytoplasm during intraluminal vesicle formation. These structures were similar in size to small viruses such as picornaviruses (McFerran et al., 1971) or parvoviruses (Cossart et al., 1975). However, it is unlikely they are infectious viruses as we observed them in DRGs derived from a mouse colony which is regularly screened to ensure they are free from infection. In support of this, EM images of infectious viruses which contain a genome are usually ’filled’ (Vijayakrishnan et al., 2020) whereas our virus-like capsid structures do not contain internal density.

This group of non-membranous compartments may be examples of endogenous capsid assemblies such as Arc. Arc proteins self-assemble into virus-like capsids to transfer mRNA between neurons (Ashley et al., 2018; Pastuzyn et al., 2018) and are key regulators of synaptic plasticity and long-term memory formation (Plath et al., 2006; Shepherd and Bear, 2011). However, purified recombinant rat Arc capsids have mean diameter of 32 nm *in vitro* (Pastuzyn et al., 2018) and Drosophila Arc capsids have diameter 37 nm (Erlendsson et al., 2020) which are both larger than the particles we observe. The virus-like capsids may alternatively correspond to an as-yet unidentified protein compartment. Bacteria and archaea contain nanocompartments called encapsulins which are also capsid-like structures (Giessen et al., 2019). EM studies of these have shown these are 20-40 nm in diameter and bind proteins in their core for storage and protection (Lau et al., 2018; McHugh et al., 2014; Sigmund et al., 2018; Sutter et al., 2008). In our tomograms we see small, globular density in the lumen of some virus-like capsids (Figure 5B,D), suggesting they may also be storage compartments. On average we found 1.4 virus-like capsids per micron of axon, suggesting they are an abundant component that are distributed throughout axons.

## Outlook

Previous descriptions of axonal ultrastructure using cryo-ET have mainly focused on synaptic regions. In this study, we characterize the axon shaft, the cellular conduit which supplies and maintains synaptic connections. Our analysis has allowed us to describe many unanticipated features including clusters of connections between the ER and microtubules, granules in the lumen of vesicles and an abundance of proteinaceous capsid-like particles in the axoplasm. This work demonstrates that new features of cells can be uncovered using cryo-ET even when they have been well studied by other imaging methods.

Our work also presents the challenge of identifying the components that make up these novel structures. A key limitation for cryo-ET imaging is the thickness of samples. Our data shows that when sufficiently thin regions are imaged, fine structures of the axon such as connections between membranes and microtubules are visible. Further acquisition and analysis of high quality cryo-ET images will enable generation of subtomogram average structures in the future, allowing the structure of these complexes to be solved and the components identified. Taken together, this study provides insight into the molecular architecture of the axoplasm.

## Materials and Methods

### DRG neuron culture on EM grids

Primary dorsal root ganglia (DRG) neuron cultures were derived from 6-8 week old mice as described in (Tolkovsky and Brelstaff, 2017). All procedures were carried out in accordance with UK Home Office regulations and licensed under the UK Animals (Scientific Procedures) Act of 1986 following local ethical approval. Spines from male and female, wild-type or transgenic GFP-RFP-LC3 mice (gift from David Rubinsztein, CIMR) we obtained. Cervical and thoracic ganglia were isolated from bisected spines and submerged in ice-cold HBSS (Thermo Fisher) supplemented with 20 mM HEPES pH 7.4 (Sigma) (HBSS+H). Ganglia from 2-4 mice were pooled and the excess connecting axon was removed. After transfer into a 15 mL Falcon, the ganglia were washed two times in 2 mL HBSS+H, then resuspended in 1 mL HBSS warmed to 37°C and supplemented with 15 µL 20mg/mL collagenage type IV. After incubation for 1 hour at 37°C, 5% CO_2_, 1 mL warm HBSS supplemented with 100 µL 2.5% Trypsin was added. After 15 mins, the enzymatic digestion was quenched by adding 5 mL warm plating media (Neurobasal media (Thermo Fisher) supplemented with 1 x B-27 (Thermo Fisher), 2 mM L-Glutamine (Thermo Fisher), 20 mM HEPES pH 7.4, 100 U/mL penicillin, 100 U/mL streptomycin and 5% FBS (Thermo Fisher). Cells were washed twice by in 2 mL plating media and resuspended in 1 mL plating media before being sequentially triturated using a 1000 µL then 200 µL pipette. Depending on the number of spines dissected, the cell suspension was loaded onto 2-4 ice-cold 3 mL 15% BSA-DMEM cushions and the cell bodies were pelleted for 8 min at 300g (4°C). After discarding the supernatant, the DRG cell bodies were gently resuspend in 200 µL plating media pre-warmed to 37°C.

Quantifoil R3.5/1 Au 200 mesh grids (Quantifoil Micro Tools) were prepared by coating with a thin layer of homemade continuous carbon and then cleaned for 40 s in a Nano Clean Plasma cleaner Model 1070 (Fischione) at 70% power in a 9:1 mixture of Argon and Oxygen gas. Soon after treatment, each grid was transferred into a microwell of an Ibidi µ-slide 2 well co-culture dish (Ibidi). In a laminar flow hood, 0.1 mg/mL sterile poly-L-lysine (Sigma-Aldrich) diluted in water was added and incubated for 4-6 hours at 37°C. After washing twice in water, 0.01mg/mL sterile laminin (Sigma-Aldrich) was added for overnight incubation. Immediately prior to plating, the laminin solution was removed and grids were washed twice in water then once in plating media.

Cells were diluted to 60,000 cells/mL and 40 µL added to each microwell and placed in a 37°C incubator at 5% CO_2_. After 2 hours, the macrowells were topped up to 700 µL with maintenance media consisting of Neurobasal media, 1 x B-27, 2 mM L-Glutamine, 20 mM HEPES pH 7.4, 100 U/mL penicillin, 100 U/mL streptomycin and 100 ng/µL NGF (PeproTech). The next day, half of the media was replaced with maintenance media supplemented with 40 µM UfdU (Sigma-Aldrich) to suppress proliferation of mitotic cells. Cultures were maintained for 3-7 days before fluorescent imaging or vitrification.

### Fluorescent imaging of acidic vesicles

Imaging was performed on neurons grown for 3 days *in vitro* (DIV), at 37°C, in 5% CO_2_ and all media were pre-warmed to 37°C before labelling or washing. For fluorescent labelling of acidic vesicles, 50 nM LysoTracker dye (Thermo Fisher) diluted in maintenance media was added to the culture dish containing EM grids. After 5 mins, the grid was transferred to a new culture chamber containing fresh culture media and inverted so the cell-plated side was facing the objective lens. If the grid was sufficiently flat, the focus position of a grid square could be maintained during image acquisition. Images were acquired on an Andor Revolution spinning disk using a far-red filter set every 500 ms for 2 min, with 50 ms exposure for each image. ImageJ (Schindelin et al., 2012) was used for projection of maximum intensity in Z (over time) and kymograph generation.

### Cryo-EM sample preparation

After verification of axon growth by light microscopy, neurons grown on grids were vitrified by manual blotting within a Vitrobot MkII (Thermo Fisher). The Vitrobot chamber was set to 37°C, 100% humidity with blotting arms disabled. 4 x 0.7 cm strips of Whatman No. 1 filter paper were folded so the final 1 cm was at a 90° angle to the length of the strip and kept in a warm, humidified chamber for at least 20 mins before blotting. Each grid was lifted from the culture and 4ul fresh culture medium containing 5 nm BSA-Au (BBI Solutions) diluted 1:6 was added before loading into the Vitrobot. Grids were blotted from the back for 3-5 s using the 1 x 0.7 cm folded section of the filter paper strips held in forceps before plunging into liquid ethane.

### Cryo-ET data acquisition

Tomograms were acquired using a Titan Krios microscope (Thermo Fisher Scientific) operated at 300 kV and equipped with an energy filter operated with slit width 20 eV. SerialEM (Mastronarde, 2005) was used to acquire tilt series between ±60° in 2° increments. At each tilt angle, a movie containing 10 frames during exposure of 2 e^-^/Å^2^ was acquired on a K2 camera (Gatan). Dataset 1 was acquired on GFP-RFP-LC3 neurons at DIV 4 on MRC-LMB Krios 3 at nominal pixel size 3.44 Å/pix using a zero-centered tilt scheme from -30° to +60° then -30° to -30° (30/32 tomograms acquired were reconstructed). Dataset 2 was acquired on wild-type neurons at DIV 7 on eBIC Krios 1 at nominal pixel size 2.75 Å/pix using a dose-symmetric tilt scheme (Hagen et al., 2017) (39/43 tomograms acquired were reconstructed). Defocus for each tomogram was set to between -3 and -6 µm underfocus.

### Tomogram reconstruction and visualization

The raw images were gain- and motion-corrected using the alignframes IMOD (Kremer et al., 1996) program. During this process, per-frame dose weighting was performed, taking into account the cumulative dose during the tilt series. Tilt series with errors during data collection were not processed and only those with images for tilts ±52° were retained. Tomogram alignment and reconstruction was performed using the ETomo interface of IMOD. Although gold fiducials were added to the samples, data collection was not limited to regions with abundant fiducials. Where fewer than 5 fiducials where present, patch tracking was performed. All tomograms were reconstructed using weighted back projection. They were then binned by 4 and filtered for visualization in MATLAB, using the Wiener-like deconvolution filter implemented in WARP (Tegunov and Cramer, 2019) (https://github.com/dtegunov/tom_deconv). The resulting bin4, deconvolved tomograms were inspected in IMOD using the slicer window and features annotated by generation of IMOD models.

### Characterization of filaments and vesicles

To determine if filaments have regular structure along their length, we performed Fourier analysis of individual filaments after their alignment in the IMOD Slicer window. The position of reflections was determined in ImageJ after contrast adjustment.

Measurement of filament length was determined using IMOD models. The ends of microtubules and ’thick’ filaments were rarely observed within the imaged volumes and likely have lengths longer than the 1.2 µm field of view. Both ends of actin and ’thin’ filaments were frequently observed and their length was measured. This is referred to as visible length as we can not rule out that the filaments do extend for longer than this. The diameters of filaments, vesicles and non-membrane bound compartments as well as the length of the ER-to-microtubule connections were also measured in IMOD. For all quantification, mean ± standard deviation is given.

For quantification of the difference in vault wall thickness compared to lipid bilayers, we first calculated the average membrane thickness in five tomograms containing a vault. In IMOD, the width of five lipid bilayers were measured in each tomogram. We then measured the wall thickness of each vault on three different places and compared each vault wall thickness to the average lipid bilayer thickness in that tomogram to give the fold change in vault width compared to membranes.

### Subtomogram averaging of ’thick’ filaments

’Thick’ filaments were modelled and picked using a similar workflow to that described for microtubule averaging in (Foster et al., 2021). 565 ’thick’ filaments subtomograms (from 13 filaments) were extracted from 7 bin4 tomograms with 24 nm particle spacing. After randomization of the in-plane angle, the particles were averaged using the TOM and AV3 packages (Förster and Hegerl, 2007; Nickell et al., 2005) operated via the subTOM software package (as described in (Leneva et al., 2020), available at https://github.com/DustinMorado/subTOM/releases/tag/v1.1.4).

### Secondary structure prediction of ER-resident proteins

Secondary structure prediction of mouse Sec61β (Q9CQS8), REEP1 (Q8BGH4) and p180 (RRBP1, Q99PL5) was performed using PSIPred (Jones, 1999) and was depicted if the confidence score was above 0.7 for more than 5 residues. Disorder prediction was performed using MobiDB Lite (Necci et al., 2017). Helices previously reported to be transmembrane were confirmed by TMHMM (Krogh et al., 2001) and coiled coil propensity predicted using COILS (Lupas et al., 1991).

## Author contributions

H.E.F and A.P.C conceived the project. H.E.F. performed data acquisition and analysis. C.V.S aided data interpretation and together with H.E.F prepared figures. A.P.C guided the project. All authors wrote the manuscript.

## ACKNOWLEDGEMENTS

We thank the MRC-LMB Electron Microscopy Facility for access and support of electron microscopy sample preparation and data collection. We acknowledge Diamond for access and support of the Cryo-EM facilities at the UK national electron bio-imaging centre (proposal EM17434-85), funded by the Wellcome Trust, MRC and BBSRC. We also thank the Light Microscopy, Scientific Computing and Biological Services facilities at the MRC-LMB for their help and support. We thank D. Morado for training and support in tomography data acquisiton and analysis. We acknowledge F. Siddiqi and D. Rubinsztein for the gift of the GFP-RFP-LC3 spines. We thank members of the Carter lab for critical reading of the manuscript.

---

**Supplementary Figure S1.**
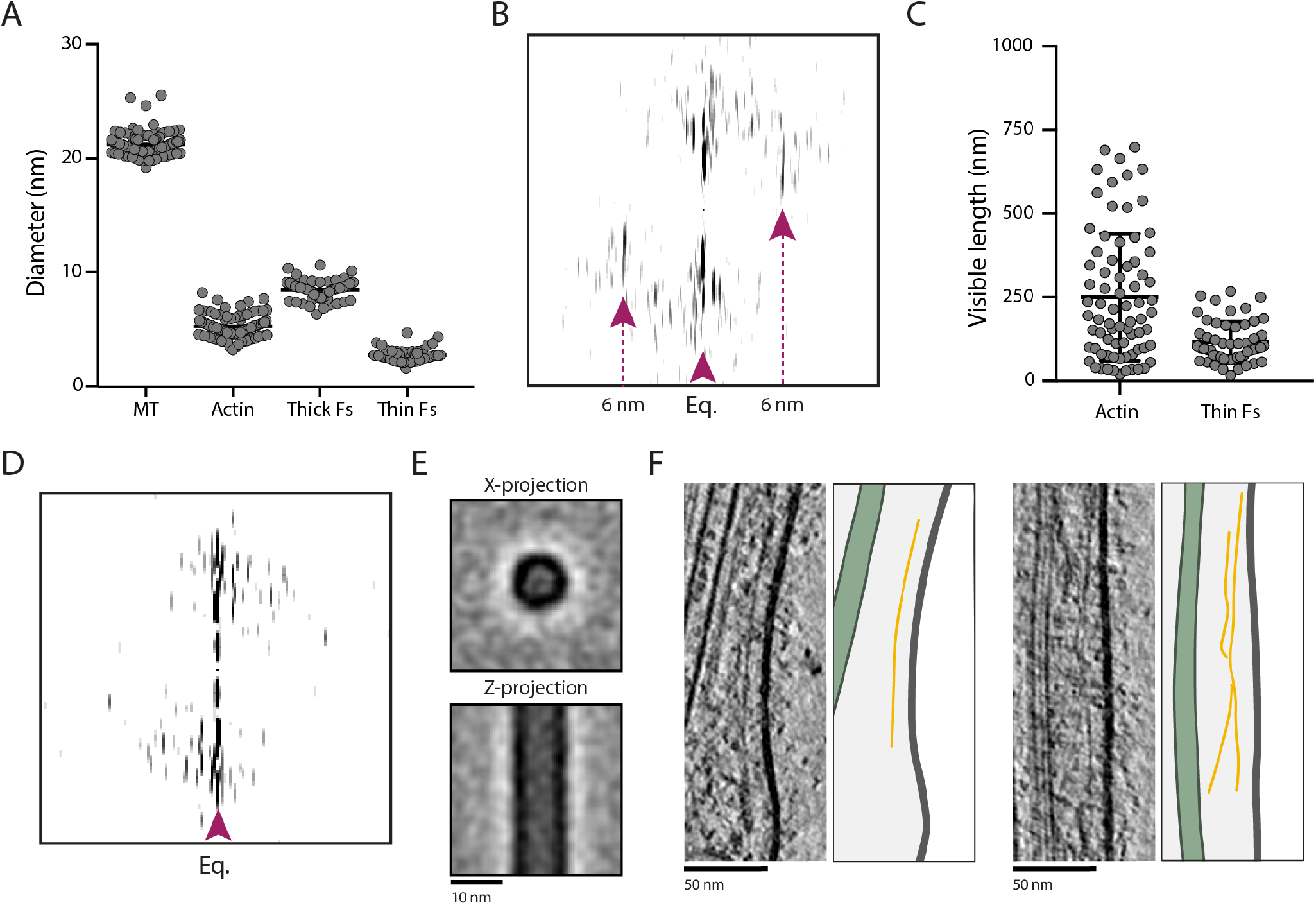
Properties of axonal filaments. A) The diameter of microtubules (MT, 21.2±1.0 nm, n=113), actin (6.1±1.1 nm, n=91), ’thick’ filaments (thick Fs, 8.5±0.8 nm, n=62) and ’thin’ filaments (thin Fs, 2.8±0.6 nm, n=48) measured from 2D slices of cryo-electron tomograms. B) Fourier transform of actin filament showing characteristic 6 nm layer lines. Position of equator (Eq.) and layer lines are indicated with purple arrows. C) The visible length of actin (251.3±190.2 nm, n=75) and thin Fs (116.2±62.4 nm, n=53) within cryo-electron tomograms. D) Fourier transform of a thick F showing no clear reflections along the length of the filament. Position of equator (Eq.) is indicated with purple arrowhead. E) Subtomogram average structure of thick Fs show they are tubular. 5.5 nm slices of the average projected from the top (X) or side (Z) are shown. F) Slices through tomograms showing thin Fs.

**Supplementary Figure S2.**
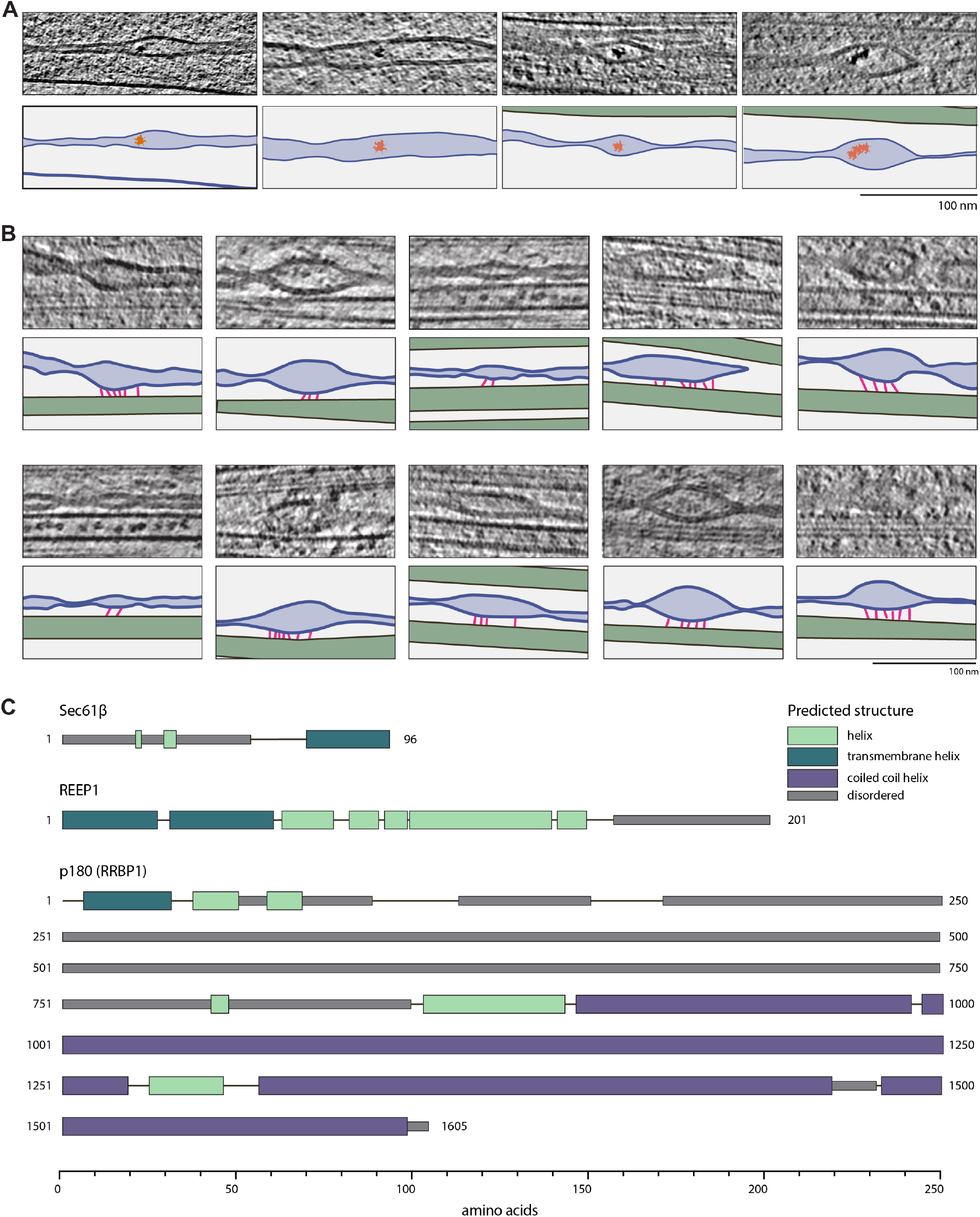
Gallery of ER-microtubule tether sites and structure prediction of candidate factors. A) Slices through tomograms showing regions of ER with luminal deposits. In the model below each example, microtubules are green, ER membrane is blue, ER lumen is light blue and deposits are orange. B) Gallery of ER-microtubule contact sites. Bottom panels are colored as in A. C) Schematic representations of secondary structure predictions for ER-resident transmembrane (TM) proteins which bind microtubules. PSIPRED 4.0, MobiDB Lite, TMHMM and COILS were used for secondary structure, disorder, transmembrane helix and coiled coil helix prediction respectively.

**Supplementary Figure S3.**
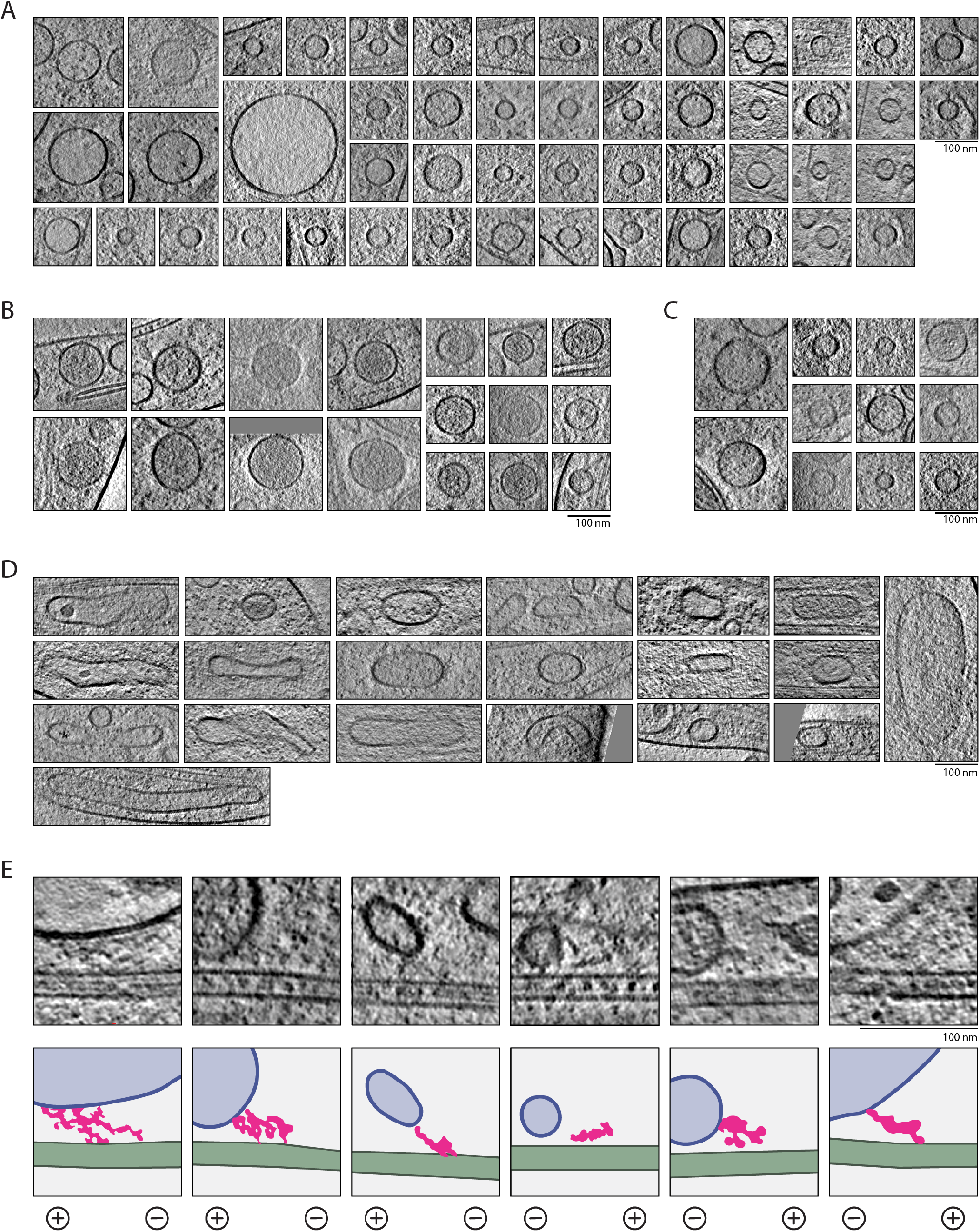
Gallery of uni-lamellar membrane compartments and connections to microtubules. A) Slices through tomograms showing the morphology of light-lumen spherical vesicles. (B) As in A, for dark-lumen spherical vesicles. C) As in A, for spherical vesicles with ambiguous lumen density. D) As in A, for elongated vesicles. E) Slices through tomograms showing microtubules close to membrane-bound compartments where connecting density is visible. In the models below each example, microtubules are green, membranes are blue, lumen is light blue and connecting density is pink. The microtubule orientation is indicated below each example.

**Supplementary Figure S4.**
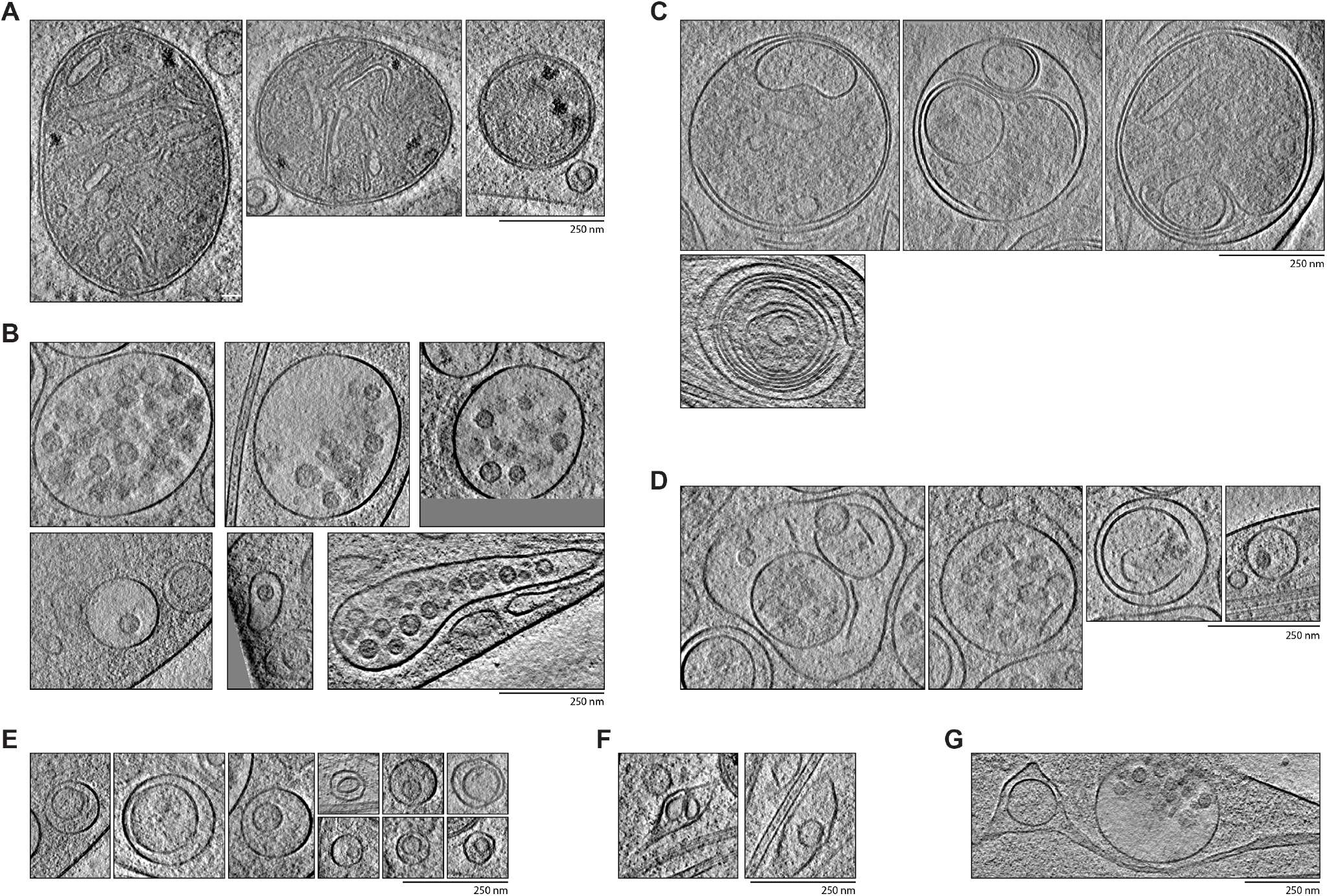
Gallery of multilamellar compartments. A) Slices through tomograms showing examples of mitochondria. B) As in A, for multivesicular bodies. C-D) Slices through tomograms showing multimembrane compartments with broken membranes in their lumen. Examples in C are more electron dense and appear more similar to autophagosomes while those in D and less electron dense and are more similar to endolysosomes. E-G) Slices through tomograms showing compartments with unclear identity. Examples in E) contain large, internal vesicles enclosed by a single spherical membrane. Examples in F) are within elongated compartments and G) shows a vesicle within the ER lumen.

**Supplementary Figure S5.**
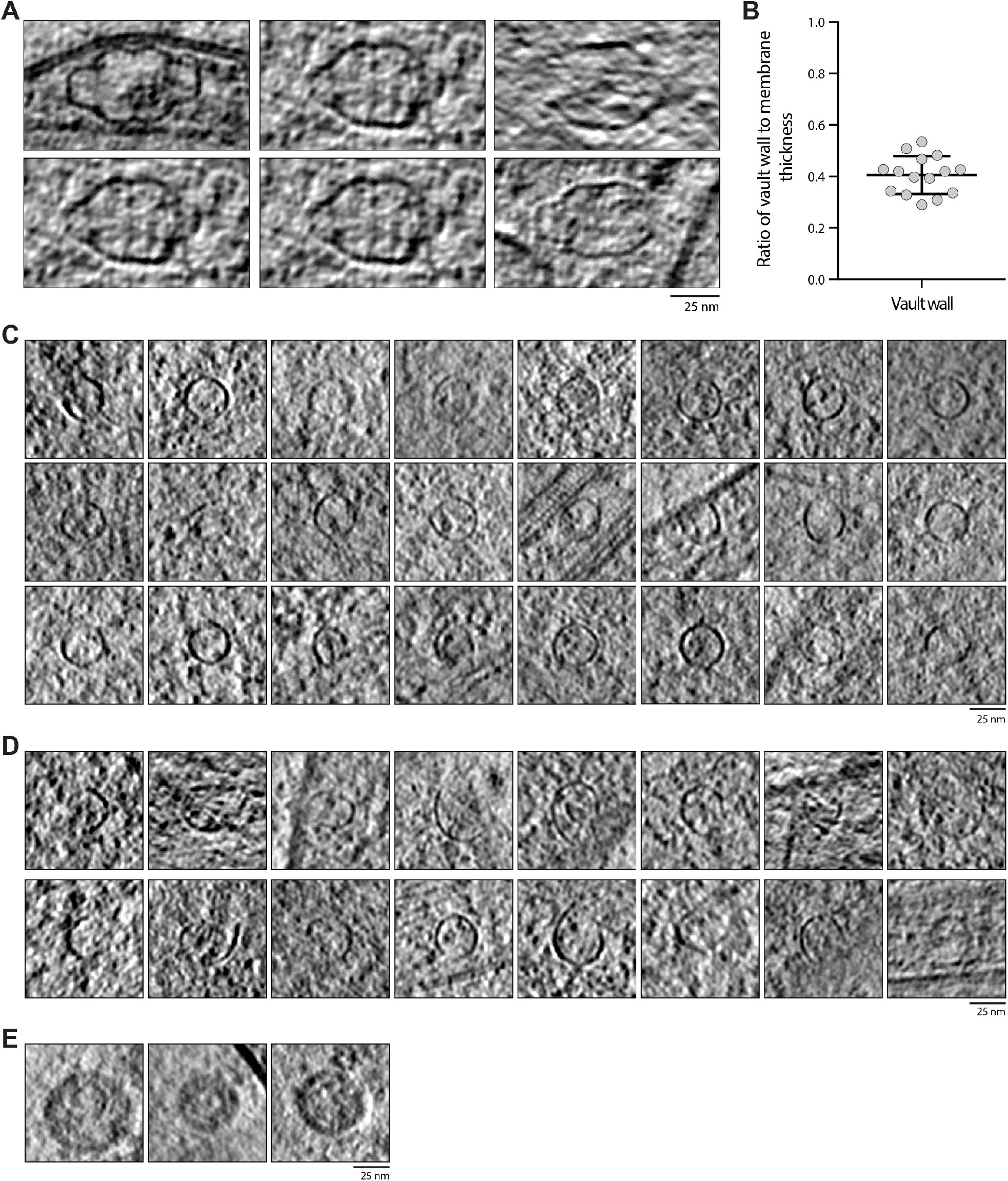
Gallery of axonal non-membrane bound compartments. A) Slices through tomograms showing examples of vaults. B) Graph showing the vault wall is less than half the width of the membrane. The relative fold change in width for each vault (n=15) wall measurement was compared to the average membrane width calculated from 5 measurements in the same tomogram as the vault. C-E) Examples of virus-like capsids which are intact (C), open (D) or within intraluminal vesicles (E).

